# Influence of environmental temperature on mouth-form plasticity in *Pristionchus pacificus* acts through *daf-11*-dependent cGMP signaling

**DOI:** 10.1101/2021.05.22.445254

**Authors:** Maša Lenuzzi, Hanh Witte, Metta Riebesell, Cristian Rödelspereger, Ray L. Hong, Ralf J. Sommer

**Affiliations:** Max-Planck Institute for Developmental Biology, Department for Integrative Evolutionary Biology, Max-Planck Ring 9, 72076 Tübingen, Germany; California State University, Northridge, Department of Biology, 18111 Nordhoff St. Northridge CA, 91330, U.S.A.

**Keywords:** *Pristionchus pacificus*, developmental plasticity, developmental switch genes, guanylyl cyclases, cyclic nucleotide-gated channels, *Caenorhabditis elegans*

## Abstract

Mouth-form plasticity in the nematode *Pristionchus pacificus* has become a powerful system to identify the genetic and molecular mechanisms associated with phenotypic (developmental) plasticity. In particular, the identification of developmental switch genes that can sense environmental stimuli and reprogram developmental processes has confirmed long-standing evolutionary theory. Together with the associated gene regulatory networks, these developmental switch genes have been important to show that plasticity is consistent with the Modern Synthesis of evolution. However, how these genes are involved in the direct sensing of the environment, or if the switch genes act downstream of another, primary environmental sensing mechanism, remains currently unknown. Here, we study the influence of environmental temperature on mouth-form plasticity. Using forward and reverse genetic technology including CRISPR/Cas9, we show that mutations in the guanylyl cyclase *Ppa-daf-11*, the *Ppa-daf-25*/AnkMy2 and the cyclic nucleotide-gated channel *Ppa-tax-2* eliminate the response to elevated temperatures. Together, our study indicates that DAF-11, DAF-25 and TAX-2 have been co-opted for environmental sensing during mouth-form plasticity regulation in *P. pacificus*. This work suggests that developmental switch genes integrate environmental signals including perception by cGMP signaling.

## Introduction

Distinct environmental conditions can shape development, resulting in different, sometimes even alternative phenotypes (Stearns, 1989; Pigliucci, 2001). Such ‘developmental (phenotypic) plasticity’ has been argued to facilitate evolutionary novelty and the origin of complex traits (West-Eberhard, 2003), a claim that has subsequently been confirmed in different study systems including vertebrates and invertebrates (Levis et al., 2018; Susoy et al., 2015; Parker and Brisson, 2019; van der Burg et al., 2020). As such, developmental plasticity is of importance for both developmental biology and evolutionary biology and highlights the significance of environmental factors (Sommer, 2020). However, the mechanistic basis of developmental plasticity was long limited to hormones (Suzuki and Nijhout, 2006). Only in the last decade did studies begin to identify associated developmental switches and gene regulatory networks (GRN). To this end, studies in genetically identical organisms that reproduce by asexual reproduction or self-fertilization (hermaphrodites) have provided important molecular insight, which includes among others, work in the water flea *Daphnia*, the pea aphid *Acyrthosiphon pisum*, hymenopterans with their elaborate caste systems and the self-fertilizing nematodes *Caenorhabditis elegans* and *Pristionchus pacificus*.

To validate the significance of plasticity for development and evolution, two lines of molecular evidence must be taken into account. First, what is the genetic and molecular basis of plastic trait development and how does it differ from ‘hard-wired’ developmental processes that are insensitive to environmental input? Second, how is the environmental information sensed and subsequently transmitted resulting in the formation of different phenotypes? The nematode *Pristionchus pacificus* has emerged as one promising model organism for the investigation of phenotypic plasticity with forward and reverse genetic tools (Sommer et al., 2017). *P. pacificus* shares with *C. elegans* many life history traits, including hermaphroditic reproduction and a short life cycle that is completed after four days when grown on standard nematode-growth-medium (NGM) agar plates with *E. coli* OP50 as food source at 20 °C (Sommer et al., 1996). *C. elegans* and *P. pacificus* are members of different nematode families, the Rhabditidae and Diplogastridae, respectively, and are thought to have separated roughly 100 million years ago (Prabh et al., 2018; Werner et al., 2018a). Given its hermaphroditic mode of reproduction, *P. pacificus* is amenable to genetic manipulation including forward genetic mutagenesis, CRISPR knockouts and engineering and DNA-mediated transformation (Sommer and Sternberg, 1996; Nakayama et al., 2020; Han et al., 2020). The *P. pacificus* genome was originally sequenced in 2008 (Dieterich et al., 2008) with more recent single molecule re-sequencing and transcriptomic efforts, which indicated that the roughly 170 Mb genome contains approximately 28,000 genes (Rödelsperger et al., 2017, 2019; Athanasouli et al., 2020).

One of the most prominent differences between *P. pacificus* and *C. elegans* is that *P. pacificus* exhibits plasticity of its feeding structures with two alternative mouth-forms that contain teeth-like denticles (Bento et al., 2010). These structures represent novel and complex traits unknown from most nematodes. Specifically, adult *P. pacificus* express either a narrow *stenostomatous* (St) morph with a single flint-like dorsal tooth, or a wide *eurystomatous* (Eu) morph, which has a claw-like dorsal tooth and an enlarged right subventral tooth (Figure 1A). Importantly, juveniles and adult St animals are strict bacterial feeders, whereas adult Eu animals can additionally kill other nematodes and subsequently feed on them (Figure 1A). Thus, morphological mouth-form plasticity results in alternative behaviors and feeding strategies. Several aspects of the mouth-form dimorphism have made it a powerful system to investigate the genetics of phenotypic plasticity. First, the discrete polyphenism allows categorical phenotyping without intermediate forms. Second, the hermaphroditic mode of reproduction mitigates any concern that genetic polymorphisms contribute to experimentally observed phenotypes. Third, hermaphroditism allows for the isolation of mutants that affect mouth-form ratios (Ragsdale et al., 2013). Finally, environmental influence can easily be manipulated under laboratory conditions. For example, when wild type worms of the *P. pacificus* reference strain PS312 are cultured on NGM agar plates they exhibit predominantly the Eu morph (>90% Eu), whereas they are only 10% Eu when grown in liquid culture (Werner et al., 2017). Thus, *P. pacificus* mouth-form plasticity can be genetically and environmentally manipulated allowing for a detailed analysis of associated molecular mechanisms.

**Fig. 1:**
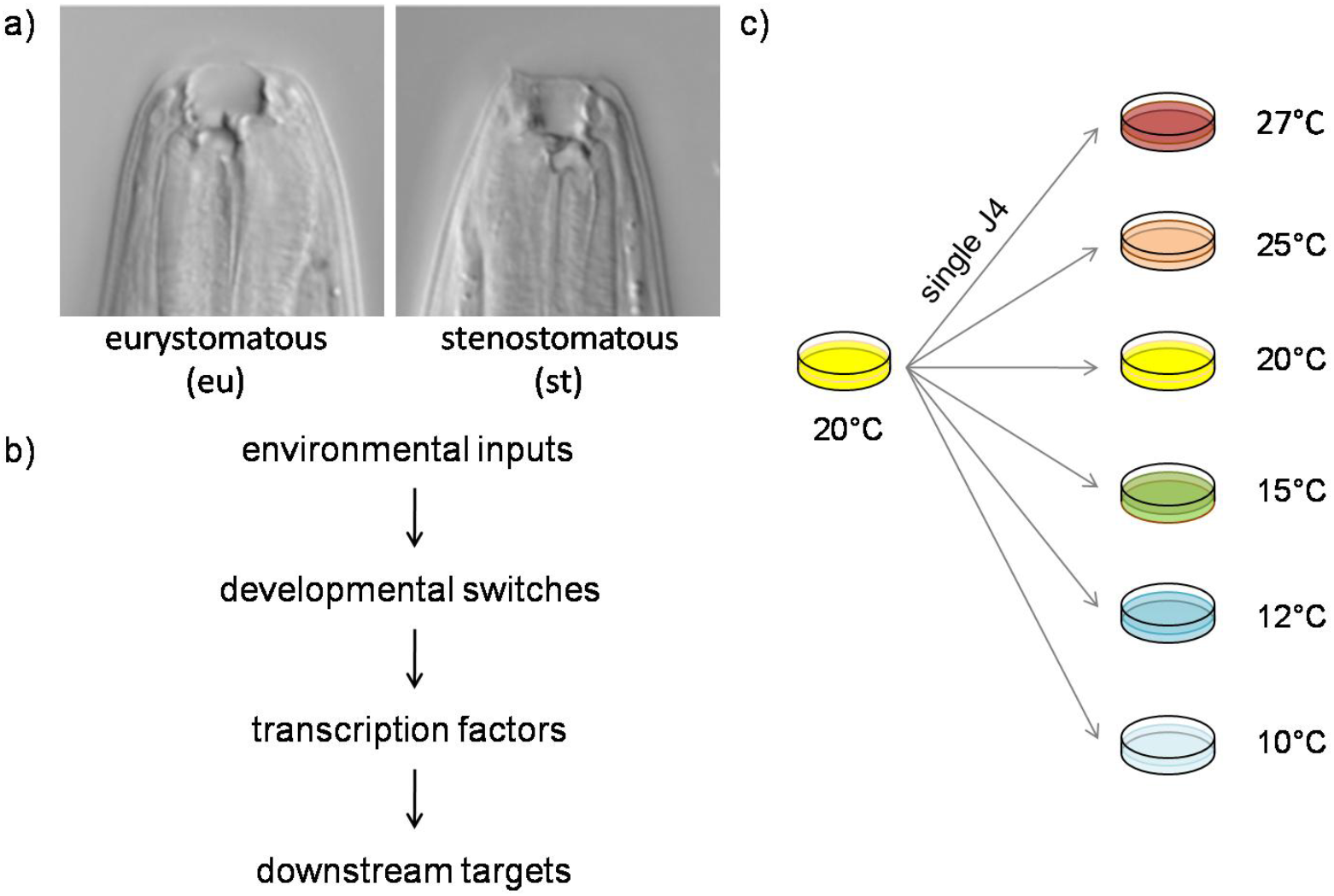
*P. pacificus* mouth-form polyphenism, conceptual perspective of mouth-form regulation and study design. A) *P. pacificus* forms two alternative mouth forms. The eurystomatous (Eu) morph is a potential predator of other nematodes, has a wide buccal cavity and two teeth, one of which is visible in this plane of focus. The stenostomatous (St) morph is a strict bacterial feeder, has a narrow mouth and single dorsal tooth. B) Conceptual perspective of the mouth form gene regulatory network. Environmental signals are sensed by the developing animal through i) a developmental switch that can reprogram development, ii) downstream transcription factors and their iii) final targets. C) Study design for testing the effect of environmental temperature. Single J4 larval animals from a standard culture at 20 °C are transferred to the test temperature and their F1 progeny are phenotyped.

What is the genetic basis of mouth-form plasticity? A series of forward and reverse genetic investigations in the last decade has gradually resulted in the discovery of a mouth-form gene regulatory network (MF-GRN) that includes i) developmental switches, ii) at least two transcription factors of the nuclear-hormone-receptor family, and iii) their downstream targets (Figure 1B) (Bui et al., 2018; Kieninger et al., 2016; Namdeo et al., 2018; Ragsdale et al., 2013; Serobyan et al., 2016; Sieriebriennikov et al., 2018, 2020, Bui and Ragsdale, 2019; Casasa et al., 2020).

The first gene to be identified came from *eu*rystomatous-form-*d*efective *eud-1* mutants, which remained St animals under all culturing conditions tested (Ragsdale et al., 2013). Most importantly, *eud-1* is dose sensitive and thus represents a developmental switch. This finding has confirmed long-standing predictions about developmental switches as key elements of developmental plasticity, which allow environmental sensing and developmental reprogramming (West-Eberhard, 2003; Ragsdale et al., 2013; Sommer, 2020). Interestingly, *eud-1* resides in a conserved ‘multi-gene locus’ and is surrounded by tandem-duplicated genes (*nag-1* and *nag-2*), which promote the St morph (Sieriebriennikov et al., 2018). While *nag-1* and *nag-2* mutants are all-Eu, *eud-1* mutant animals are all-St with *eud-1* being epistatic (Sieriebriennikov et al., 2018). All three genes, *eud-1, nag-1* and *nag-2* are expressed in different sensory neurons. The identification of switch genes that govern intra-generational plasticity is important to confirm that plasticity is indeed consistent with the Modern Synthesis of evolution (Baugh and Day, 2020). However, it remained unclear if these genes are involved in the direct sensing of the environment or if they act downstream of another, primary environmental sensing mechanism.

Here, we study the influence of environmental temperature on mouth-form plasticity (Figure 1C). We show that mouth-form plasticity in many *P. pacificus* isolates is strongly sensitive to the temperature of the environment. We focus on the isolate RSA635 from La Réunion island that is 70% Eu at 20°C, but only 20% Eu at 27 °C. Using a combination of forward and reverse genetic approaches, we found that cGMP signaling is involved in temperature perception, which in turn regulates mouth-form plasticity. First, mutations in the guanylyl cyclase *Ppa-daf-11* and the *Ppa-daf-25*/AnkMy2 eliminated the response to elevated temperatures as revealed in forward genetic screens. Second, reverse genetic approaches showed that the cyclic nucleotide-gated channel *Ppa-tax-2*, but not the Forkhead-type transcription factor *Ppa-daf-16* required for temperature perception. Together, our study indicates that the DAF-11, DAF-25 and TAX-2 orthologs have been co-opted for environmental sensing during mouth-form plasticity regulation in *P. pacificus*. This work suggests that developmental switch genes integrate environmental signals that at least in part are perceived by cGMP signaling and potentially other primary signaling systems.

## Results

### *P. pacificus* shows strain-specific temperature sensitivity of mouth-form plasticity

To study the molecular mechanisms of environmental sensing during mouth-form plasticity regulation in *P. pacificus*, we investigated the influence of temperature as an abiotic factor for several reasons. First, temperature is known to be involved in the regulation of plastic traits in a diversity of animals and plants (Pigliucci, 2001). Second, the molecular mechanisms of temperature perception in development and physiology have been intensively studied in several model organisms, including comprehensive investigations in *C. elegans* (for review see, Goodman and Sengupta, 2019). Finally, *P. pacificus* exhibits higher temperature tolerance than *C. elegans* with several tested wild isolates being capable of reproducing at 30 °C (Leaver et al., 2016). Therefore, we first compared the mouth-form ratios of the wild type strain *P. pacificus* PS312 (RS2333) grown at 10 °C, 15 °C, 20 °C, 25 °C and 27°C on standard NGM agar plates (Figure 1C). At most temperatures we observed more than 80% Eu animals, which means the observed temperature effect was not very strong (Figure2). Only at temperatures below 15 °C, when worms grow very slowly, we found less than 50% of animals having the Eu mouth form. These findings suggest that temperature has a limited effect on mouth-form ratio in *P. pacificus* RS2333, which might result from an original adaptation of this genotype or from domestication processes. Note that RS2333 is a direct derivate of the original PS312 strain that has been isolated in Pasadena, California in 1988 and has been in laboratory culture ever since (Sommer et al., 1996). Next, we investigated 10 natural isolates of *P. pacificus* that cover the large genetic diversity of *P. pacificus*, including several isolates from the tropical island La Réunion in the Indian ocean (Rödelsperger et al., 2014, McGaughran et al., 2016). Most noticeably, none of these strains has undergone a long period of domestication in the laboratory and most of them have been frozen in the first 20 generations after isolation. Indeed, testing the same set of temperatures in these strains revealed a strong sensitivity of mouth-form plasticity to environmental temperature (Figure2). Specifically, many strains show a bell-like curve with the highest percentage of Eu animals observed at 20 °C, such as in RSA622, RSA635 and RSA645. In other strains, the highest Eu ratios are found at 15 °C or 25 °C culture conditions. Additionally, some strains, like RS5160, do not form more than 50% Eu animals at any temperature. We conclude that in the large majority of *P. pacificus* strains environmental temperature has an influence on mouth-form plasticity, largely in a strain-specific pattern, and that this effect can likely be subjected to genetic investigations.

### Forward genetic screens identify temperature sensing-deficient mutants

To initiate genetic analysis of temperature perception of *P. pacificus* mouth-form plasticity, we selected strain RSA635 with its bell-shaped temperature response curve. At 20 °C, around 70% of the animals express the Eu mouth form, whereas only 20% are Eu at the two extreme temperatures 10 °C and 27°C (Figure 2, Table 1A). We performed EMS mutagenesis with RSA635 J4 juveniles using the standard *P. pacificus* mutagenesis protocol (Aurilio and Srinivasan, 2015). Subsequently, we examined the F3 progeny of 2,400 F2 animals for high Eu mouth-form ratios after culturing the F3 generation at 27 °C. We isolated 13 mutant candidates with >70% Eu animals at 27 °C. After backcrossing, we recovered 11 mutant lines with elevated Eu mouth-form ratios. Several of the mutant strains, however, showed high variability in mouth-form ratios. Three of these 11 mutant lines, *tu715, tu719* and *tu724*, were 100% Eu at 27 °C (Table 1B). This phenotype is similar to the phenotype of mutations in the *nag-1* and *nag-2* genes in the *P. pacificus* wild type, which form the multi-gene switch locus (Sieriebriennikov et al., 2018).

**Table 1:** Mouth-form frequency in wild type and mutant lines.

|  | Mutagen | Genotype | Molecular lesion | %Eu 20°C<br>(N) | % Eu 27°C<br>(N) |
| --- | --- | --- | --- | --- | --- |
| A | n.a. | RSA635, WT | n.a. | 71(248) | 21(201) |
| B | EMS | *Ppa-nag-1(tu715)* | nonsense | 100(30) | 100(99) |
|  | EMS | *Ppa-nag-1(tu719)* | non-synonymous | 100(30) | 100(69) |
|  | EMS | *Ppa-nag-1(tu724)* | non-synonymous | 100(30) | 100(96) |
| C | EMS | *Ppa-daf-25(tu716)* | non-synonymous | 90(30) | 99(128) |
|  | CRISPR | *Ppa-daf-25(tu1516)* | 11 bp insertion | 100(90) | 93(90) |
|  | CRISPR | *Ppa-daf-25(tu1517)* | 8 bp deletion | 98(90) | 91(90) |
| D | EMS | *Ppa-daf-11(tu722)* | nonsense | 93(30) | 93(86) |
|  | CRISPR | *Ppa-daf-11(tu1438)* | 2bp deletion | 78(86) | 87(63) |
|  | CRISPR | *Ppa-daf-11(tu1439)* | 2 bp deletion | 87(90) | 80(67) |
|  | CRISPR | *Ppa-daf-11(tu1440)* | 4bp insertion | 79(90) | 84(90) |
| E | CRISPR | *Ppa-daf-16(tu1514)* | 5bp insertion | 43(85) | 22(32) |
|  | CRISPR | *Ppa-daf-16(tu1515)* | 11 bp insertion | 51(84) | 13(44) |
| F | CRISPR | *Ppa-daf-16(tu1515),*<br>*Ppa-daf-11(tu1440)* | n.a. | 43(85) | 22(32) |
| G | CRISPR | *Ppa-tax-2(tu1291)* | 20 bp insertion | 95(60) | 88(42) |
|  | CRISPR | *Ppa-tax-2(tu1292)* | 32 bp insertion | 82(51) | 60(10) |

All lines are generated in the RSA635 background. EMS, Ethylene-methylsulfonate; CRISPR, mutations generated by CRISPR/Cas9 knockouts. n.a., not applicable

**Fig. 2:**
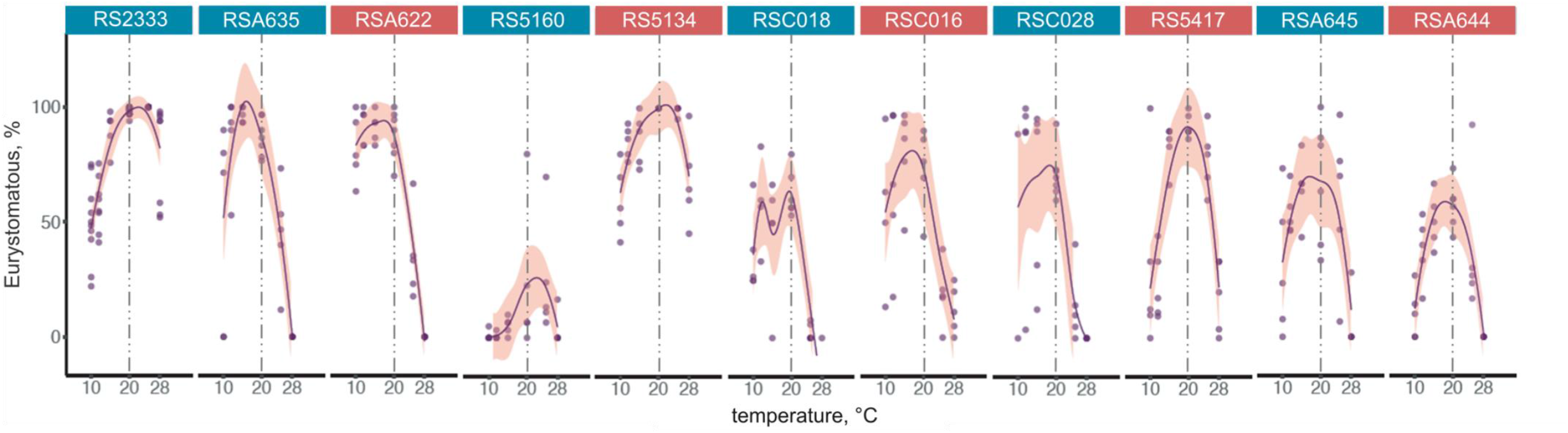
Natural variation of the reaction norm of *P. pacificus* RS2333, a direct derivate of the original isolate PS312 from Pasadena (California, US) and 10 wild isolates. High temperature tolerant strains are labeled in blue, low-temperature tolerant strains in red. Vertical line indicates the mouth-form frequency at 20 °C. Eu percentages for each line and temperature were plotted using geom_smooth function of ggplot2 R package.

To test whether these mutant lines affect the mouth-form switch, we phenotyped them at 20 °C and performed whole genome sequencing (WGS) to identify potential causal mutations. Indeed, all three mutant lines were 100% Eu at 20 °C and thus, the effect on mouth-form ratio in these mutant lines is temperature-independent (Table 1B). Additionally, WGS identified mutations in *nag-1* in all three mutant lines. Specifically, *tu715* has a nonsense mutation, whereas *tu719* and *tu724* result in nonsynonymous changes in *Ppa-nag-1* (Table 1B). Together, these experiments indicate that mutants with altered temperature perception during plasticity regulation can be isolated in *P. pacificus* RSA635. Furthermore, these findings suggest that the multi-gene locus controlling mouth-form switching is conserved among strains and acts in a temperature-independent manner.

### *tu716* is caused by a mutation in *daf-25*, an ortholog of mammalian Ankmy2

From the remaining eight mutant lines with elevated Eu mouth-form ratios at 27 °C, we selected two lines for further analysis. These mutants, *tu716* and *tu722*, showed the most consistent phenotype over multiple generations. For example, *tu716* is 99% Eu at 27 °C and 90% Eu at 20 °C, a phenotype that is less extreme than the one observed in the three *Ppa-nag-1* alleles (Table 1C). We mapped *tu716* through a modified bulk segregant analysis regime (see Materials and Methods for details). In short, we sequenced 23 mutant animals after backcrossing with RSA635 wild type. We generated Tn5 single-worm libraries for all 23 mutant animals, includinga similar number of non-mutant control animals from the same cross. After sequencing, we performed SNP enrichment in the mutant batch and identified a potential mutation in the *P. pacificus* ortholog of *daf-25*. In *C. elegans, daf-25* mutants show a temperature-sensitive *da*uer *f*ormation*-c*onstitutive (Daf-c) phenotype (Jensen et al., 2010). *daf-25* is localized in cilia of sensory neurons and encodes the ortholog of the mammalian Ankmy2, a MYND domain protein of unknown function (Jensen et al., 2010).

*tu716* exhibits a missense mutation in the predicted protein pocket of *Ppa-*DAF-25 resulting in a V > E amino acid change (Table 1C). To verify that the temperature sensing defect of *tu716* is indeed due to the mutation in *Ppa-daf-25*, we generated additional alleles by CRISPR knockout, a method that has been applied successfully in previous studies (Moreno et al., 2017). We isolated two CRISPR-induced mutants in *Ppa-daf-25* using a guide RNA in the fourth exon, close to the site of the mutation in the original *tu716*allele. The new alleles, *Ppa-daf-25 (tu1516)* and *Ppa-daf-25 (tu1517)*, have an 11 bp insertion and an 8 bp deletion, respectively (Table 1C). Both alleles are highly Eu at 27 °C, similar to *Ppa-daf-25 (tu716)* (Table 1C). However, none of the *Ppa-daf-25* alleles displayed the Daf-c phenotype. We conclude that *Ppa-daf-25* is involved in temperature perception of mouth-form plasticity. These results are of particular interest because in *C. elegans*, DAF-25 is known to be required for the proper localization of the guanyl cyclase DAF-11 in cGMP signaling in cilia (Jensen et al., 2010).

### A *daf-11*/guanylyl cyclase mutant is high temperature insensitive

Next, we used a similar mapping strategy for the second mutant with a strong temperature-sensing defect. *tu722* is 93% Eu at 27 °C and 93% Eu at 20 °C (Table 1). Using the same bulk segregant analysis regime we found a mutation in *Ppa-daf-11* in exon 18, resulting in a premature stop codon (Table 1D). This observation would be consistent with the known interaction of *daf-25* and *daf-11* in *C. elegans* and with the results described above. We used a CRISPR knockout approach to confirm this observation by generating additional mutants in *Ppa-daf-11*. Specifically, we used one sgRNA to induce mutations close to the N-terminus of the gene in exon 3 and a second sgRNA in exon 18, the same exon that harbors the *tu722* mutation. The sgRNA in exon 3 resulted in five mutant F2 lines with deletions in *Ppa-daf-11*. However, all these lines were sterile so that no homozygous mutant line could be generated. This finding suggests that *loss-of-function* or strong *reduction-of-function* alleles of *Ppa-daf-11* cannot be kept as homozygous mutant lines. In contrast, we isolated three viable alleles with mutations in exon 18. Specifically, *Ppa-daf-11 (tu1438)* and *Ppa-daf-11 (tu1439)* have two bp deletions and *Ppa-daf-11 (tu1440)* has a four bp insertion, all of which result in premature stop codons. All three of these mutant lines show strongly elevated Eu mouth-form frequencies at 27 °C, similar to the original *Ppa-daf-11 (tu722)* allele (Table 1D). Thus, the guanylyl cyclase *Ppa-daf-11* is involved in temperature perception during mouth-form plasticity regulation. The fact that *Ppa-daf-11* and *Ppa-daf-25* have similar phenotypes is consistent with the previously described role of *Cel*-DAF-25 in the proper localization of *Cel*-DAF-11 to cilia in *C. elegans* (Jensen et al., 2010). Together, these findings allow two conclusions. First, cGMP signaling involving *Ppa-daf-11* and *Ppa-daf-25* is required for temperature sensitivity of mouth-form plasticity in *P. pacificus*. Second, the *daf-11/daf-25* module has been co-opted during nematode evolution for regulating the influence of temperature on mouth-form plasticity in *P. pacificus*.

### *Ppa-daf-11* is expressed in multiple amphid neurons

In *C. elegans, daf-11* mutants are Daf-c, forming dauer larvae in the absence of dauer-inducing conditions (Riddle et al., 1981; Thomas et al., 1993). Additionally, *Cel-daf-11* mutants exhibit defects in several chemosensory responses, including the attraction to volatile odorants such as isoamyl alcohol (Vowels and Thomas, 1994). Consistent with most of these phenotypes, *Cel-daf-11* is expressed in the five pairs of amphid neurons ASI, ASJ, ASK, AWB and AWC (Birnby et al., 2000). To study if the co-option of *Ppa-daf-11* for temperature perception during mouth-form plasticity regulation involved novel expression patterns, we generated *Ppa-daf-11p::rfp* reporter lines. We used a 2 kb promoter fragment and obtained two transgenic lines *tu*Ex329 and *tu*Ex330 with 30% and nearly 100% transmission rate, respectively. *Ppa-daf-11p::rfp* is expressed in five pairs of amphid neurons (Figure3). According to the recent neuroanatomical study of the chemosensory system of *P. pacificus* (Hong et al., 2019), these cells are likely the amphid neurons AM8(ASJ), AM1(ASH), AM3(AWA), AM5(ASE) and AM4(ASK) (Figure 3). These assignments suggest similarity and divergence of *daf-11* expression in amphid neurons, a finding that is related to previous observations. For example, *Ppa-odr-7* and *Ppa-odr-3* also show partial conservation and divergence of gene expression (Hong et al., 2019). Specifically, *Ppa-daf-11* and *Ppa-odr-3* are expressed in the AWA homolog rather than the AWC-equivalent neurons. It is notable that the absence of the wing-type *C. elegans* amphid neurons (AWA, AWB, AWC) in *P. pacificus* is shared with all other nematodes that have been studied by EM reconstruction (Hong et al., 2019).

**Fig. 3:**
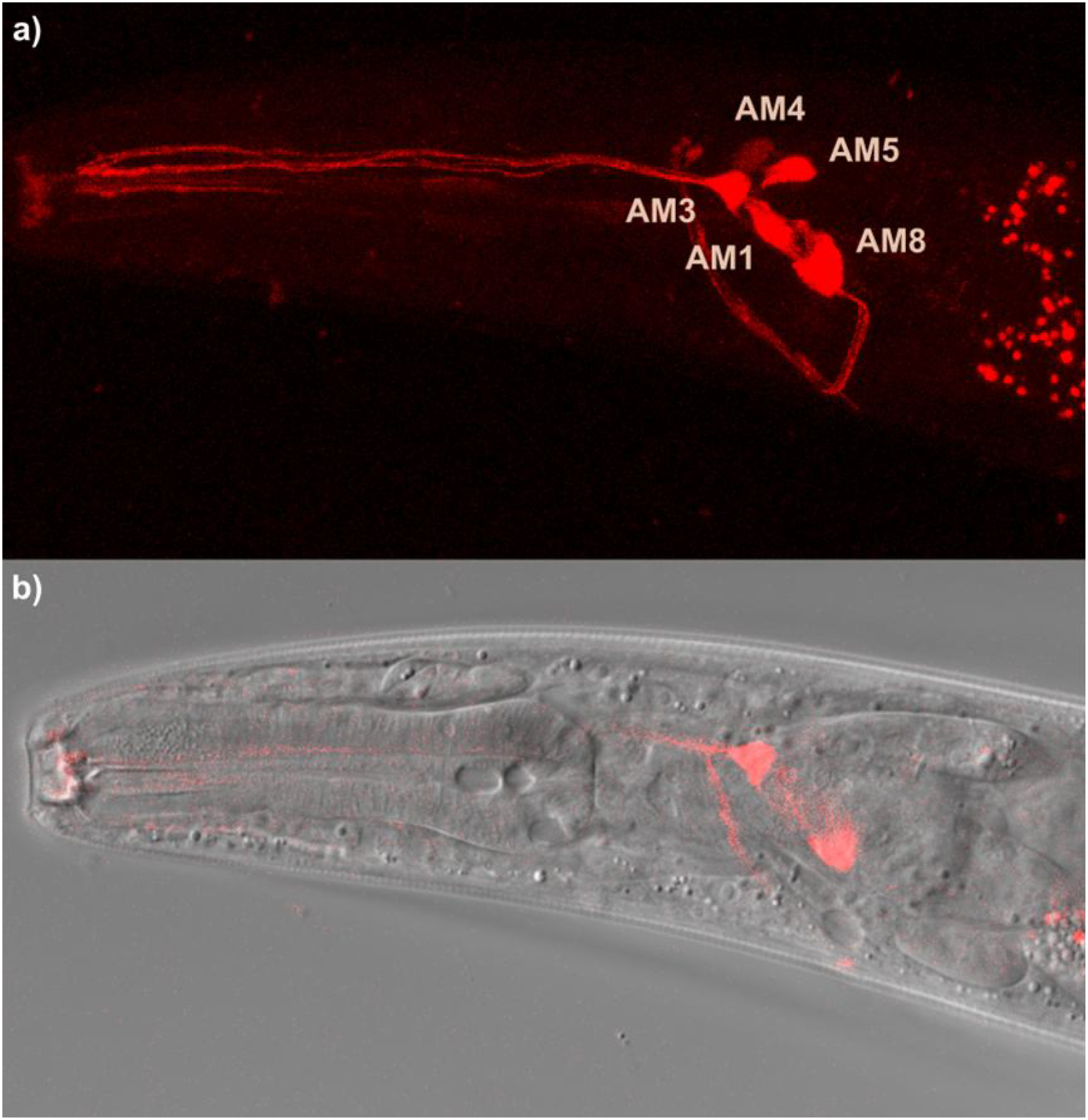
Expression pattern of *Ppa-daf-11* as revealed by *Ppa-daf-11*::TurboRFP reporter. A) Expression in the head region in a young adult in transgenic line tuEx330. The expression in the head was visible in both juvenile and adult stages (males and hermaphrodites). Z-stacks of the anterior part of adult hermaphrodites were acquired and further analyzed using maximum projection. See text for identity of amphid neurons. B) Corresponding Normaski image.

### *Ppa-tax-2* but not *Ppa-daf-16* is required for temperature sensing during mouth-form regulation

Next, we wanted to identify additional components of the gene regulatory network involved in temperature perception. We used a candidate gene approach and targeted genes through CRISPR-mediated gene knockouts. First, we selected the FOXO transcription factor *daf-16* because it is known as a terminal regulator of many developmental and physiological processes including dauer formation in *C. elegans* (Kenyon, 2010). The analysis of *Ppa-daf-16* for a potential role in mouth-form plasticity is important because previous studies did not find any mouth-form associated phenotype (Ogawa et al., 2011). While *Ppa-daf-16* mutants are *da*uer *f*ormation-*d*efective (daf-d), like in *C. elegans*, no change in mouth-form ratios was observed in these mutants. However, these mutants in the *P. pacificus* RS2333 background showed nominal temperature response (Figure 2). Therefore, we generated two novel *Ppa-daf-16* mutant alleles in the RSA635 background, *tu1514 and tu1515*. When we analyzed the mouth-form ratio of these mutantsraised at 27 °C, we did not observe any difference from RSA635 wild type animals (Table 1E). However, when cultured at 20 °C, both *Ppa-daf-16* alleles showed only 43% and 51% Eu animals (Table 1E). Given that in *C. elegans* dauer development, *daf-16* is negatively regulated by *daf-11*-dependent cGMP signaling, we made a *Ppa-daf-16 (tu1515), Ppa-daf-11 (tu1440)* double mutant. These double mutantshad a high Eu phenotype at 27 °C similar to *Ppa-daf-11* single mutants (Table 1F), suggesting that *Ppa-daf-16* is not involved in the regulation of mouth-form plasticity in *P. pacificus*.

Finally, we wanted to know if *Ppa-daf-11* and *Ppa-daf-25* function through a canonical cGMP signaling pathway, or alternatively, through a different molecular network. Work in *C. elegans* indicated that different cGMP signaling pathways involve different guanylyl cyclases that converge on the two cyclic nucleotide-gated channels *tax-2* and *tax-4* (Aoki and Mori, 2015). Therefore, we generated two*Ppa-tax-2* mutants in the RSA635 background. Indeed, *Ppa-tax-2 (tu1291)* and *Ppa-tax-2 (tu1292)* mutant animals show elevated Eu mouth-form ratios when cultured at 27 °C similar to *Ppa-daf-11* and *Ppa-daf-25* mutants (Table 1G). Thus, *Ppa-daf-11* acts through a canonical cGMP signaling pathway in temperature perception during mouth-form plasticity regulation.

## Discussion

This study presents a developmental genetic analysis to elucidate the influence of environmental temperature on mouth-form plasticity in *P. pacificus*. Our findings reveal that temperature has a strong effect on the mouth-form decision. Together with previous studies, these results indicate that *P. pacificus* decides which mouth-form to execute mainly in response to pheromones (Bose et al., 2012; Werner et al., 2018b), nutrient conditions (Werner et al., 2017) and temperature (this study). Thus, like dauer formation in *C. elegans* (Golden and Riddle, 1984) and *P. pacificus* (Ogawa et al., 2009), mouth-form plasticity is regulated by a set of biotic and abiotic factors. This study produced in two major conclusions.

First, our study indicates that temperature perception acts mainly through cGMP signaling and the *Ppa-*DAF-25/Ankmy2 – *Ppa-*DAF-11 module. In *C. elegans*, the guanylyl cyclase-encoding gene *daf-11* was isolated as a ‘group 1 dauer’ mutant with a strong Daf-c phenotype (Thomas et al., 1993). Additionally, *Cel-daf-11* mutants have defects in chemosensation, *i*.*e*. they do not respond to isoamyl alcohol (Vowels and Thomas, 1994) and were later shown to have complex defects in ASJ sensory neurons (Schackwitz et al., 1996). The localization of the guanylyl cyclase to chemosensory cilia requires the MYND/Ankmy2 domain protein DAF-25 and consequently, *C. elegans* mutants in *daf-11* and *daf-25* show similar Daf-c phenotypes (Jensen et al., 2010).

The fact that we have identified *Ppa-daf-11* and *Ppa-daf-25* in the same screen for temperature influence on mouth-form plasticity suggests that both proteins have similar and conserved functions in *P. pacificus*. Phylogenetically, mouth-form plasticity represents a more recently evolved trait than dauer formation. Specifically, mouth-form plasticity is restricted to the Diplogastridae family, whereas the potential to form dauer larvae under adverse conditions is much more widespread in nematodes. Therefore, the DAF-11/DAF-25 module has likely been co-opted to control temperature perception in mouth-form plasticity. Co-option is a well-established mechanism by which genes become integrated in the regulation of novel developmental and physiological traits (True and Carroll, 2002). Indeed, co-option was already identified as a major principle in evolutionary developmental biology when the first systematic studies of molecular comparative developmental biology were performed in insects, nematodes and vertebrates (Raff, 1996).

Second, DAF-11 is one of more than 30 guanylyl cyclases in the genomes of *C. elegans* and *P. pacificus*, many of which are 1:1 orthologous (Hong et al., 2019). In C. *elegans* a subfamily of guanylyl cyclases, *gcy-8, gcy-18* and *gcy-23*, have a major additional role in temperature sensation primarily through the amphid finger neuron AFD (Inada et al., 2006). Indeed, besides the influence of temperature on certain developmental and chemosensory processes as indicated above, *C. elegans* is also able to acclimate to its cultivation temperature. The resulting thermotaxis behavior becomes obvious on thermal gradients when worms prefer the temperature to which they have been previously adapted (for review see Aoki and Mori, 2015). Interestingly, the three AFD-specific guanylyl cyclases GCY-8, GCY-18 and GCY-23 are the primary thermosensors as indicated by ectopic expression of these genes conferring thermosensory properties to diverse cell types (Takeishi et al., 2016). The cyclic nucleotide-gated channels TAX-2 and TAX-4 act downstream of guanylyl cyclases in the perception of environmental temperature in various processes (Komatsu et al., 1996). Consequently, *tax-2* and *tax-4* mutants in *C. elegans* show abnormalities in multiple sensory behaviors with thermotaxis phenotypes similar to the *gcy-8, gcy-18, gcy-23* triple mutants, as well as dauer formation and chemotaxis phenotypes similar to *daf-11* mutants. Our candidate gene approach indicates that *Ppa-*TAX-2 is also involved in the transmission of cGMP signaling during *P. pacificus* mouth-form control. These findings suggest that a major temperature perception network consisting of *Ppa-*DAF-11, *Ppa-*DAF-25, *Ppa-*TAX-2 and presumably other factors acts as one first step in the environmental regulation of mouth-form plasticity in *P. pacificus*. We speculate that other factors control the influence of pheromones and nutrients on mouth-form plasticity. We hypothesize that all relevant environmental cues are integrated at the level of the switch network, consisting of the sulfatase EUD-1 and the N-acetyl-glucosaminidases NAG-1 and NAG-2 (Sieriebriennikov et al., 2018). However, the exact molecular mechanism of this regulatory interaction awaits future analysis. Our finding that mutations in the FORKHEAD-type transcription factor *Ppa-daf-16* do not share the mouth-form phenotypes of *Ppa-daf-11* and *Ppa-tax-2* at culture conditions at 27 °C may indicate the involvement of additional factors, which might even act independently of transcriptional regulation.

Finally, we note that we have not observed any spontaneous dauer formation in *Ppa-daf-25* and *Ppa-daf-11* mutants on culturing plates, neither at 20 °C nor at 27 °C. These observations are superficially similar to previous findings that the regulation of dauer larvae formation has undergone substantial changes in the lineages leading to *P. pacificus* and *C. elegans*. For example, *Ppa-daf-19*, the RTX master transcriptional regulator of ciliogenesis has no dauer phenotype in the RS2333 background (Moreno, Lenuzzi et al., 2018). To clarify this point further, more detailed analyses of dauer formation have to be performed using the various available dauer-induction protocols in *P. pacificus* (Markov et al., 2016; Werner et al., 2018b) - a project which is clearly outside of the scope of the current manuscript.

In conclusion, our study establishes the influence of environmental temperature on mouth-form plasticity in *P. pacificus* and reveals the co-option of cGMP signaling as one mechanism of environmental sensing. This work adds important information to the complexity of the gene regulatory network involved in the regulation of this novel and complex trait. The identification of a temperature perception network adds an important step in the elucidation of the molecular mechanisms associated with this novel trait. A proper understanding of the proximate developmental mechanisms is crucial for a full acceptance of developmental plasticity as an important mechanism for evolution.

## Acknowledgment

The study was funded by the Max Planck Society. The authors declare no conflict of interest. We thank members of the Sommer lab for discussion and assistance throughout these studies.

## Author contributions

Conceptualization, M.L., R.J.S., Methodology, M.L., H.W., M.R.; Analysis, M.L., M.R., R.H.; Resources, H.W.; Writing – Original draft, R.J.S.; Writing – Review and Editing, R.J.S., M.L., M.R., R.H.

## Materials and Methods

### Strains and mutagenesis experiments

The reference strain *P. pacificus* RS2333 from Pasadena (California, USA) and the wild isolates RS5134 (USA), RS5160 (Japan), and the La Réunion and Mauritius-derived strains RS5417, RSA622, RSA635, RSA644, RSA645, RSC016, RSC018 and RSC028 of *P. pacificus* were used for studying natural variation of mouth-form. The population genetic analysis of these strains has been described by McGaughran and co-workers (2016). All strains were maintained using standard methods (Pires daSilva, 2018).

### Mouth-form phenotyping

Mouth-form phenotyping was scored as previously described (Serobyan et al., 2013). Briefly, Eu and St individuals were distinguished by the presence or absence of a subventral tooth and a claw-like *vs*. flint-like dorsal tooth. Phenotypes were mostly consistent across temperatures, with some intermediate phenotypes on extreme temperatures as previously described (Sieriebriennikovet al., 2017). Phenotyping was performed using Zeiss Discovery V.12 and V.20 stereo microscopes and interference contrast (DIC) microscopy on a Zeiss Axioskop. For temperature screens, J4 larval stages were singled out from a mixed culture plate and transferred to a incubators (Memmert) set to the specific temperatures. The mouth-form of the next generation was screened after animals had reached adulthood (approximately 5 days on 27 °C to 3-4 weeks on 10°C).

### Mutagenesis screen for temperature insensitive mutants

We screened for temperature insensitive mutants using Ethyl methanesulfonate (EMS) in *P. pacificus* RSA635. We generated around 1,200 gametes and screened 2,400 homozygous F2 lines. F2 animals were transferred from 20°C to 27°C in the Memmert incubator on NGM plates with 200-300µl OP50. The mouth form of the F3 progeny was scored after 5-7 days.

### Bulk segregant analysis and whole-genome sequencing analysis

For bulk segregant analysis, *tu716* and *tu722* were backcrossed with wildtype animals. From each F1 backcrossed animal, four F2 worms were transferred to 27°C and the mouth-form was phenotyped after 6 days. After one generation of recovery on 20°C, individual lines were transferred again to 27°C and mouth form was scored in their progeny after 6 days. Lines with a highly Eu, or a highly St phenotype in at least three consecutive screens were used for sequencing. For that purpose, 1-3 worms were collected in lysis buffer. Whole-genome-sequencing WGS libraries were prepared using an in-house single-worm Tn5 library preparation kit. Libraries were validated using the Agilent Bioanalyzer DNA 2100 High Sensitivity DNA Assay (Agilent Technologies GmbH) and pooled prior to sequencing. Sequencing was performed on an Illumina HiSeq3000.Raw Illumina reads were aligned against the *P. pacificus* PS312 reference genome (version El Paco) with the help of the aln and sampe programs of the BWA (version 0.7.12) software suite (Li and Durbin 2010). To distinguish between mutations and variants of the different genetic backgrounds, recently generated whole genome sequencing data of the parental strain (RSA635) (McGaughran et al. 2016) was used as a control sample. To this end, differential variant calls were generated from the mpileup output of the samtools (version 0.1.18) package (Li et al. 2009) by searching for sites with at least 5X coverage where the major allele frequency was >90% but differed between the mutant and the control sample. Functional classification of differential variants was performed as described previously (Sieriebriennikov et al. 2020).

### CRISPR-Cas9 mutagenesis

We used the modified CRISPR-Cas9 protocol for *P. pacificus*as previously described (Witte et al., 2015) with hybridized target-specific CRISPR RNAs (crRNAs) and universal trans-activating CRISPR RNA (tracrRNA) obtained from Integrated DNA Technologies (Alt-R product line) for generating *Ppa-daf-25, Ppa-daf-11, Ppa-tax-2* and *Ppa-daf-16* mutants. The sgRNA and all the crRNAs targeted 20 bp upstream of protospacer adjacent motifs (PAMs). To hybridize crRNA and tracrRNA, 10 µl of the 100 µM stock of each molecule were combined, followed by a denaturation step at 95°C for 5 minutes and annealing at room temperature for another 5 minutes. Five µl of the hybridization product was combined with 1 µl of 20 µM Cas9 protein (New England Biolabs) at room temperaturefor 5 minutes. The mixture was diluted with Tris-EDTA buffer to a total volume of 25 µl and injected into the gonad of young adults. Molecular lesions were detected in F1 progeny by Sanger sequencing.

### Genetic transformation

To generate a *Ppa-daf-11*reporter construct, we amplified the upstream sequence of the in-silico identified *Ppa-daf-11* start codon to the 3’UTR of the next gene (794bp). We amplified TurboRFP and the 3’UTR of the ribosomal gene *Ppa-rpl-23* from pUC19-based plasmid stocks. All primers were obtained from Thermo Fisher Scientific. Cloning was done using Gibson Assembly® Cloning Kit (New England Biolabs). The final product was amplified as a linear construct and confirmed by Sanger sequencing. The Primers used are listed in Supplementary Table 1. For all amplification steps we used PrimeStar GxL polymerase (Takara). Prior to injection, the linear construct and genomic carrier DNA of RSA635 were incubated for two hours with PstI (New England BioLabs). The construct (60 ng/µl) was injected with the co-injection marker *egl-20*promoter::TurboRFP (10 ng/µl) and genomic DNA (1800 ng/µl) to one or both gonadal arms in early adult hermaphrodites. Images were acquired using a Leica TCS SP8 confocal system and were analyzed using ImageJ.

### Statistical analysis

All replicates of mouth-form phenotypes in mutant lines and wildtype RSA635 were compared using Fisher’s exact test. p-values were corrected using the FDR method. All mutant lines and wildtype lines were regularly observed over the course of several months to confirm the stability of the phenotype. See Supplementary materials for complete statistical analysis (Table S1-3). In figure 2, Eu percentages for each line and temperature were plotted using geom_smooth function of ggplot2 R package.

