## Supplementary material for "Influence of environmental temperature on mouth-form plasticity in *Pristionchus pacificus* acts through *daf-11*-dependent cGMP signaling": Table S1-3

**Table S1: Statistical comparison of mouth-form phenotype between wild-type RSA635 and mutant lines on 20°C.**

| **line** | **eu** | **st** | **p-value (Fisher's exact test)** | **FDR** | |
|---|---|---|---|---|---|
| wild-type | 176 | 72 | | | |
| *daf-25(tu1517)* | 89 | 1 | 3.36E-10 | 7.05E-09 | *** |
| *daf-25(tu1516)* | 70 | 0 | 2.02E-09 | 2.13E-08 | *** |
| *daf-16(tu1514)* | 52 | 63 | 4.36E-06 | 3.05E-05 | *** |
| *daf-16(tu1515)* | 55 | 59 | 5.53E-05 | 0.000184 | *** |
| *daf-11(tu1438)* | 68 | 18 | 0.1601584 | 0.174802 | ns |
| *daf-11(tu1439)* | 78 | 12 | 0.002745256 | 0.004435 | *** |
| *daf-11(tu1440)* | 71 | 19 | 0.1664776 | 0.174802 | ns |
| *tax-2(tu1291)* | 57 | 3 | 3.63E-05 | 0.000184 | *** |
| *tax-2(tu1292)* | 56 | 4 | 0.000178012 | 0.000312 | *** |
| *tax-2(tu1296)* | 42 | 9 | 0.1193128 | 0.139198 | ns |
| RS3410 | 30 | 0 | 9.66E-05 | 0.000184 | *** |
| RS3411 | 30 | 0 | 9.66E-05 | 0.000184 | *** |
| *daf-25(tu716)* | 27 | 3 | 0.02859293 | 0.037528 | * |
| RS3413 | 30 | 0 | 9.66E-05 | 0.000184 | *** |
| RS3414 | 30 | 0 | 9.66E-05 | 0.000184 | *** |
| *nag-1(tu719)* | 30 | 0 | 9.66E-05 | 0.000184 | *** |
| RS3416 | 26 | 4 | 0.08290114 | 0.102407 | ns |
| RS3599 | 27 | 3 | 0.02859293 | 0.037528 | * |
| *daf-11(tu722)* | 23 | 7 | 0.6688908 | 0.668891 | ns |
| RS3418 | 28 | 2 | 0.007713078 | 0.01157 | * |
| *nag-1(tu724)* | 30 | 0 | 9.66E-05 | 0.000184 | *** |

**Table S2: statistical comparison of mouth-form phenotype between wild-type RSA635 and mutant lines on 27°C.**

| line | eu | st | p-value (Fishers' exact test) | FDR | |
|---|---|---|---|---|---|
| wild-type | 43 | 158 | | | |
| *daf-25(tu1517)* | 58 | 7 | 4.06E-23 | 7.11E-23 | *** |
| *daf-25(tu1516)* | 61 | 4 | 9.64E-27 | 2.02E-26 | *** |
| *daf-16(tu1514)* | 10 | 52 | 4.69E-01 | 4.69E-01 | ns |
| *daf-16(tu1515)* | 7 | 46 | 0.243825 | 2.56E-01 | ns |
| *daf-11(tu1438)* | 55 | 8 | 2.66E-21 | 3.99E-21 | *** |
| *daf-11(tu1439)* | 56 | 11 | 8.33E-20 | 1.17E-19 | *** |
| *daf-11(tu1440)* | 76 | 14 | 1.44E-24 | 2.75E-24 | *** |
| *tax-2(tu1291)* | 37 | 5 | 3.11E-16 | 3.84E-16 | *** |
| *tax-2(tu1292)* | 47 | 8 | 2.81E-18 | 3.69E-18 | *** |
| *tax-2(tu1296)* | 6 | 4 | 0.011659 | 1.29E-02 | * |
| RS3410 | 37 | 6 | 1.51E-15 | 1.76E-15 | *** |
| RS3411 | 99 | 0 | 2.31E-45 | 2.43E-44 | *** |
| *daf-25(tu716)* | 127 | 1 | 4.57E-52 | 9.59E-51 | *** |
| RS3413 | 98 | 16 | 3.73E-30 | 8.69E-30 | *** |
| RS3414 | 75 | 18 | 2.46E-22 | 3.98E-22 | *** |
| *nag-1(tu719)* | 69 | 0 | 7.78E-35 | 2.33E-34 | *** |
| RS3416 | 102 | 5 | 1.11E-39 | 4.65E-39 | *** |
| RS3599 | 94 | 1 | 5.17E-42 | 2.71E-41 | *** |
| *daf-11(tu722)* | 108 | 7 | 2.26E-39 | 7.91E-39 | *** |
| RS3418 | 80 | 6 | 1.09E-31 | 2.85E-31 | *** |
| *nag-1(tu724)* | 96 | 0 | 2.20E-44 | 1.54E-43 | *** |

**Table S3: within line statistical comparison of mouth-form phenotype.**

| line.temperature | eu | st | p-value (Fisher's exact test) | FDR | |
|---|---|---|---|---|---|
| wild-type.20 | 176 | 72 | | | |
| wild-type.27 | 43 | 158 | 1.67E-26 | 1.83E-25 | *** |
| *daf-25(tu1517)*.20 | 89 | 1 | | | |
| *daf-25(tu1517)*.27 | 58 | 7 | 9.85E-03 | 2.71E-02 | * |
| *daf-25(tu1516)*.20 | 70 | 0 | | | |
| *daf-25(tu1516)*.27 | 61 | 4 | 5.12E-02 | 1.13E-01 | ns |
| *daf-16(tu1514)*.20 | 52 | 63 | | | |
| *daf-16(tu1514)*.27 | 10 | 52 | 1.21E-04 | 4.45E-04 | *** |
| *daf-16(tu1515)*.20 | 55 | 59 | | | |
| *daf-16(tu1515)*.27 | 7 | 46 | 1.17E-05 | 6.46E-05 | *** |
| *daf-11(tu1438)*.20 | 68 | 18 | | | |
| *daf-11(tu1438)*.27 | 55 | 8 | 2.75E-01 | 3.36E-01 | ns |
| *daf-11(tu1439)*.20 | 78 | 12 | | | |
| *daf-11(tu1439)*.27 | 56 | 11 | 6.51E-01 | 6.51E-01 | ns |
| *daf-11(tu1440)*.20 | 71 | 19 | | | |
| *daf-11(tu1440)*.27 | 76 | 14 | 4.41E-01 | 4.85E-01 | ns |
| *tax-2(tu1291)*.20 | 57 | 3 | | | |
| *tax-2(tu1291)*.27 | 37 | 5 | 2.69E-01 | 3.36E-01 | ns |
| *tax-2(tu1292)*.20 | 56 | 4 | | | |
| *tax-2(tu1292)*.27 | 47 | 8 | 2.26E-01 | 3.36E-01 | ns |
| *tax-2(tu1296)*.20 | 42 | 9 | | | |
| *tax-2(tu1296)*.27 | 6 | 4 | 1.98E-01 | 3.36E-01 | ns |
